# NF-κB inhibition in dendritic cells pre-treated with sulforaphane-conjugates induces immunotolerance

**DOI:** 10.1101/2024.11.27.625615

**Authors:** Camila Leiva-Castro, Ana M. Múnera-Rodríguez, Macarena Martínez-Bailén, Ana T. Carmona, Soledad López-Enríquez, Francisca Palomares

## Abstract

Sulforaphane (SFN) has notable health benefits but faces challenges due to poor solubility and delivery. This study explores SFN glycoconjugates’ effects on LPS-induced inflammation in human dendritic cells (DCs), aiming to enhance therapeutic potential against inflammatory diseases. Monovalent SFN-glycoconjugates with mannose (SFNMan) and fucose (SFNFuc) were developed and tested for their anti-inflammatory and immune-modulatory properties in DCs from healthy donors under chronic LPS exposure.

Our results revealed that carbohydrate-functionalized SFN improves solubility and effectiveness in suppressing inflammation by targeting the p65 NF-κB pathway, without affecting MAPK signaling. SFN-glycoconjugates induce a tolerogenic immune response, characterized by increased IL-10 production and enhanced regulatory T- and B-cell proliferation. Notably, these effects surpass those of p65 NF-κB inhibition alone, highlighting a distinct and potent regulatory mechanism independent of MAPK pathways.

These findings demonstrate the promise of SFN-glycoconjugates as innovative therapeutic agents for inflammatory diseases, offering enhanced anti-inflammatory and immunomodulatory effects through improved delivery and targeted molecular pathways.

## INTRODUCTION

Inflammation is a complex physiological process that plays a crucial role in the organism’s immune response to various stimuli, ranging from the presence of pathogens to tissue damage and exposure to toxins (L. Chen et al., 2018). In this context, sulforaphane (SFN), a bioactive compound found in cruciferous vegetables such as broccoli and cauliflower, has emerged as a promising candidate for modulating inflammation and its pathological consequences (Fernandez-Prades et al., 2023). SFN has been observed to exert anti-inflammatory effects by influencing key cellular signaling pathways, including the nuclear factor kappa B (NF-κB) pathways, which triggers cellular defense mechanisms, and mitogen-activated protein kinases (MAPKs), suggesting its therapeutic potential in a variety of inflammatory disorders (Deng et al., 2017; Mahn & Castillo, 2021). Indeed, our group has recently described that SFN pretreatment may induce a regulatory response by inhibiting NF-κB and MAPK signaling pathways in an inflammatory environment, along with inducing a regulatory pattern (increased IL-10-producing Treg cells) in human dendritic cells (DCs) (Fernandez-Prades et al., 2023; Munera-Rodriguez, Leiva-Castro, Sobrino, Lopez-Enriquez, & Palomares, 2024).

NF-κB is a central regulator of inflammation, controlling the expression of genes involved in immune responses. Under normal conditions, NF-κB remains inactive in the cytoplasm, sequestered by inhibitory proteins called IκBs (inhibitor of NF-κB) (L. Chen et al., 2018; Lawrence, 2009). SFN has been observed to inhibit the release of inflammatory mediators such as thymus and activation-regulated chemokine (TARC), eotaxin-1, and vascular cell adhesion molecule-1(VCAM-1) in cytokine-stimulated human corneal fibroblasts by blocking MAPK, NF-κB, and signal transducer and activator of transcription 6 (STAT6) pathways (Yang et al., 2022).

MAPKs represent a group of proteins that regulate cellular responses to diverse triggers, encompassing osmotic stress, growth factors, and inflammatory agents like interleukin (IL)-1, TNF-α (tumor necrosis factor α), and IL-6 (Burkhard & Shapiro, 2010; L. Chen et al., 2018). Dysregulation of MAPK pathways is implicated in diseases such as Alzheimer’s, Parkinson’s, and cancer (Kim & Choi, 2010). SFN’s modulation of MAPK signaling pathways may extend beyond inflammation to influence other cellular processes involved in disease pathogenesis (Zhang et al., 2022), (Saha, Buttari, Profumo, Tucci, & Saso, 2021).

Despite the promising anti-inflammatory properties demonstrated by SFN (Mahn & Castillo, 2021), its translation into clinical application has encountered challenges due to its inherent limitations in bioavailability and stability. Although SFN demonstrates relatively higher bioavailability compared to some other nutraceuticals (Caponio et al., 2022), its precise mechanism of absorption and immune cell penetration remain inadequately understood. This limited understanding hampers effective administration of SFN and its targeted delivery to specific sites of action within the body (Wang et al., 2020). Consequently, there exists a critical imperative to investigate and develop alternative strategies aimed at augmenting the delivery mechanisms and therapeutic efficacy of SFN.

In response to the challenges posed by SFN’s low bioavailability and stability, glycoconjugates (Kjaerup et al., 2014; Ramos-Soriano & Rojo, 2021) have emerged as a promising strategy to enhance its clinical efficacy in treating inflammatory and related diseases. Glycoconjugates are being actively investigated for their potential applications in vaccine development due to their ability to influence the immune response, specifically through the regulation of allergen-specific effects cells, which involves the activation of DCs (Sirvent et al., 2016).

It has been proposed that SFN-glycoconjugates, functionalized with carbohydrates such as mannose or fucose (SFNMan and/or SFNFuc), could significantly enhance and finely tune the interaction of SFN with immune cells, including DCs, thereby augmenting its therapeutic efficacy (Kjaerup et al., 2014; van Liempt et al., 2006). This innovative strategy not only aims to optimize the beneficial effects of SFN but also holds considerable promise in mitigating various inflammation-related diseases, including allergic pathologies (Rodriguez et al., 2019). By enhancing SFN’s stability and solubility through conjugation with specific carbohydrates, SFN-Man/Fuc-glycoconjugates offer a multifaceted approach to combatting inflammatory diseases, potentially leading to improved clinical outcomes and patient well-being, as observed with other glycoconjugates (Gringhuis, den Dunnen, Litjens, van der Vlist, & Geijtenbeek, 2009), (Palomares et al., 2022).

Compared to alternative administration methods, such as nanoparticles (Zambrano, Bustos, & Mahn, 2019), SFN-glycoconjugates present a range of advantages. Firstly, glycoconjugates exhibit greater stability compared to the free form of the compound, facilitating more effective interactions with immune cells (Palomares et al., 2022). Additionally, glycoconjugates can be designed to enhance the selectivity and specificity of SFN toward specific target cells, thereby reducing systemic exposure and minimizing unwanted side effects (Ribeiro-Viana et al., 2012). These advantages are particularly relevant in the context of chronic inflammatory diseases, where prolonged and targeted drug administration is essential to achieve optimal therapeutic outcomes (Palomares et al., 2019; Rodriguez et al., 2019).

In this article, we will explore the potential of SFN-glycoconjugates as a promising molecular therapy to improve the clinical efficacy of SFN in the treatment of inflammatory disorders and other related diseases. By gaining a deeper understanding of the molecular mechanisms underlying SFN-glycoconjugates and their ability to modulate inflammatory response, this research aims to develop more effective and targeted molecular therapeutic approaches to address a wide range of inflammatory and autoimmune diseases.

## RESULTS

### SFN-glycoconjugates did not produce cellular cytotoxicity

SFN-glycoconjugates were engineered to enhance certain physicochemical properties of SFN (Figure SI1A), such as stability and bioavailability, without increasing cellular toxicity. To assess this, moDCs were incubated for 48 hours with various concentrations of SFN and its glycoconjugates. Our results indicate that glycosylation of SFN increased its usable concentration in *in vitro* studies compared to SFN alone. No cytotoxic effects were observed when moDCs were stimulated with 10 μM SFN; however, higher concentrations of SFN compromised cell viability. Similarly, no cytotoxic effects were observed with the SFNMan and SFNFuc glycoconjugates at any of the concentrations tested (Figure SI1B).

### SFN-glycoconjugates modulate the p65 NF-κB but not MAPK signaling pathway

The protective effects of SFN-glycoconjugates were investigated to elucidate their molecular mechanisms in mitigating LPS-induced inflammation, focusing on the p65 NF-κB and MAPK signaling pathways, which play pivotal roles in inflammatory responses.

The SFN-glycoconjugates did not induce changes in p65 NF-κB levels compared to immature DCs. However, our results revealed that LPS instigates p65 NF-κB activation, a process significantly attenuated by SFN, SFN-glycoconjugates, and the specific p65 NF-κB blocker, MG132. Pretreatment with SFN, SFN-glycoconjugates, and MG132 for 1 hour prior to LPS exposure significantly reduced LPS-induced p65 NF-κB expression compared to LPS-pretreated moDCs (Figure 1A).

**Figure 1.**
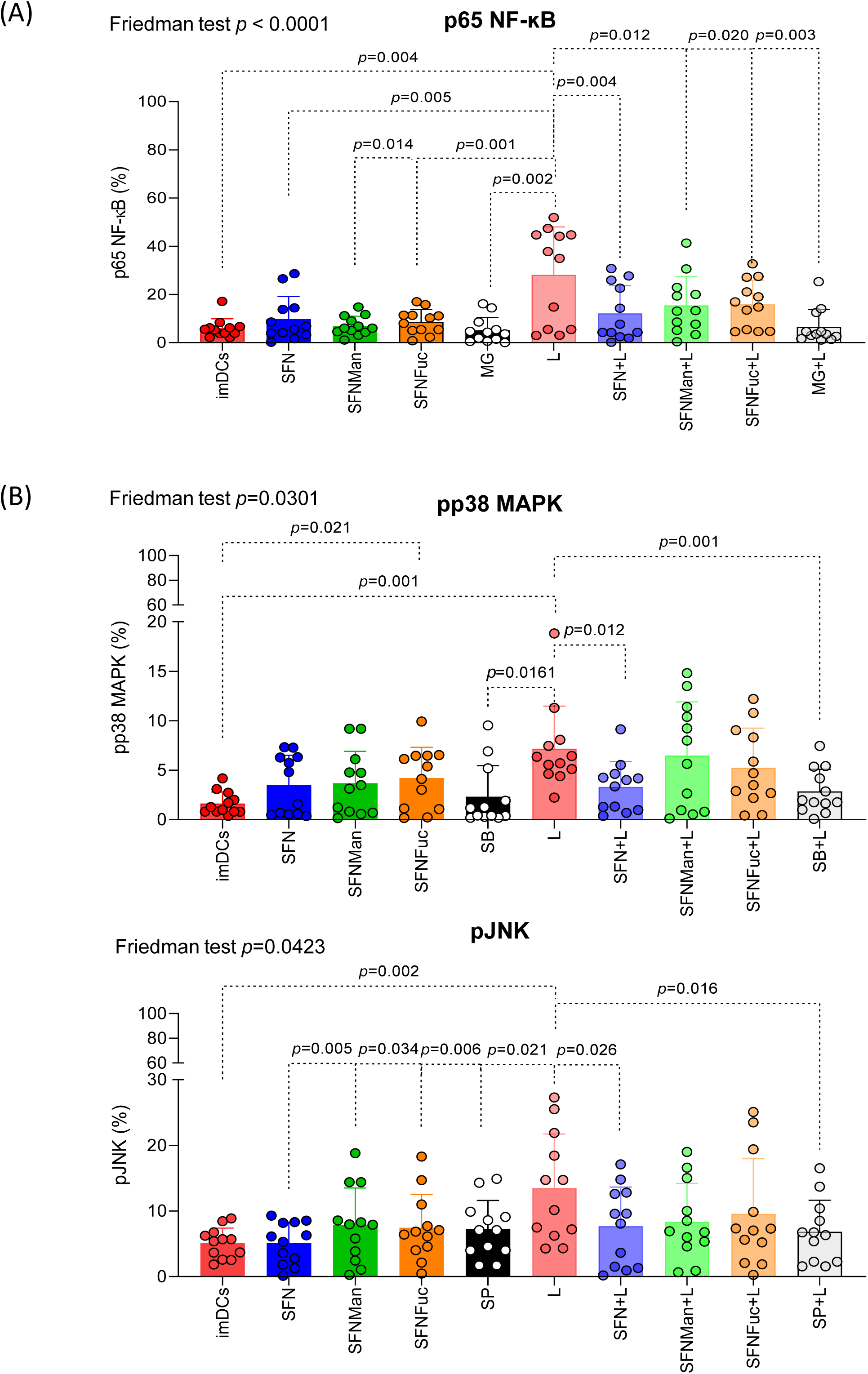
Modulation of p65 NF-κB and MAP kinase signaling pathway expression. **(A)** The bars and symbols represent the mean (SD) of the percentages of pp65 NF-κB expression, and **(B)** of different MAPK expression, for each experimental condition (N=12). imDC: immature dendritic cells. L: Lipopolysaccharide, LPS. SFN: Sulforaphane. SFNMan and SFNFuc: SFN-glycoconjugates with mannose or fucose. S+L: SFN incubated for 1 hour followed by LPS stimulation. SFNMan+ and SFNFuc+L: SFN-glycoconjugates incubated for 1 hour followed by LPS stimulation. MG: MG132, inhibitor of p65 NFkB. MG+L: MG132 incubated for 1 hour followed by LPS stimulation. SB: SB203580, blocker of p38 MAPK and SP: SP600125 as a JNK inhibitor. SB+ and SP+L: the different blockers incubated for 1 hour followed by LPS stimulation. The Friedman test was used to detect differences in related samples across multiple comparisons, indicating significant p-values. The Wilcoxon test was used for pairwise comparisons of related samples, indicating the exact p values.

Previous studies by our group demonstrated that SFN pretreatment in moDCs involved the p38 MAPK and JNK signaling pathways in modulating immune responses towards a regulatory pattern(Munera-Rodriguez et al., 2024). Based on these findings, we aimed to analyze whether SFN-glycoconjugates could regulate these signaling pathways. Results showed that LPS stimulation induced the phosphorylation of p38 MAPK (pp38 MAPK) and JNK (pJNK), with pronounced upregulation compared to immature DCs, SFN, and their respective blockers (SB203580 and SP600125). However, SFN-glycoconjugates significantly reduced pJNK levels compared to LPS-treated moDCs (Figure 1B).

Additionally, moDCs treated with SFN for 1 hour followed by LPS stimulation (S+L) demonstrated significantly reduced activation of the MAPK and JNK signaling pathways compared to LPS-treated moDCs, resembling results obtained with SB203580+L and SP600125+L (Figure 1B). Conversely, no significant differences were noted in the inhibition of the p38 MAPK and JNK pathways in SFN-glycoconjugates-pretreated moDCs challenged with LPS. These findings suggest that the chemical nature of the SFN-glycoconjugates lacks the capacity to regulate the MAPK pathways.

### SFN-glycoconjugates induce changes in the moDC maturational status

To investigate the immunomodulatory impact of SFN-glycoconjugates on moDCs, we assessed their influence on the expression of key surface regulatory molecules, specifically, CD80, CD83, CD86, PD-L1, and HLA-DR. Initially, moDCs were treated with SFN and SFN-glycoconjugates. Neither SFN nor its glycoconjugates induced changes in the expression of any markers compared to the immature state. However, a significant increase in the percentage of HLA-DR was observed in the presence of SFNFuc. In contrast, stimulation with LPS upregulated the expression of all maturation markers compared to immature DCs (Figure 2).

**Figure 2.**
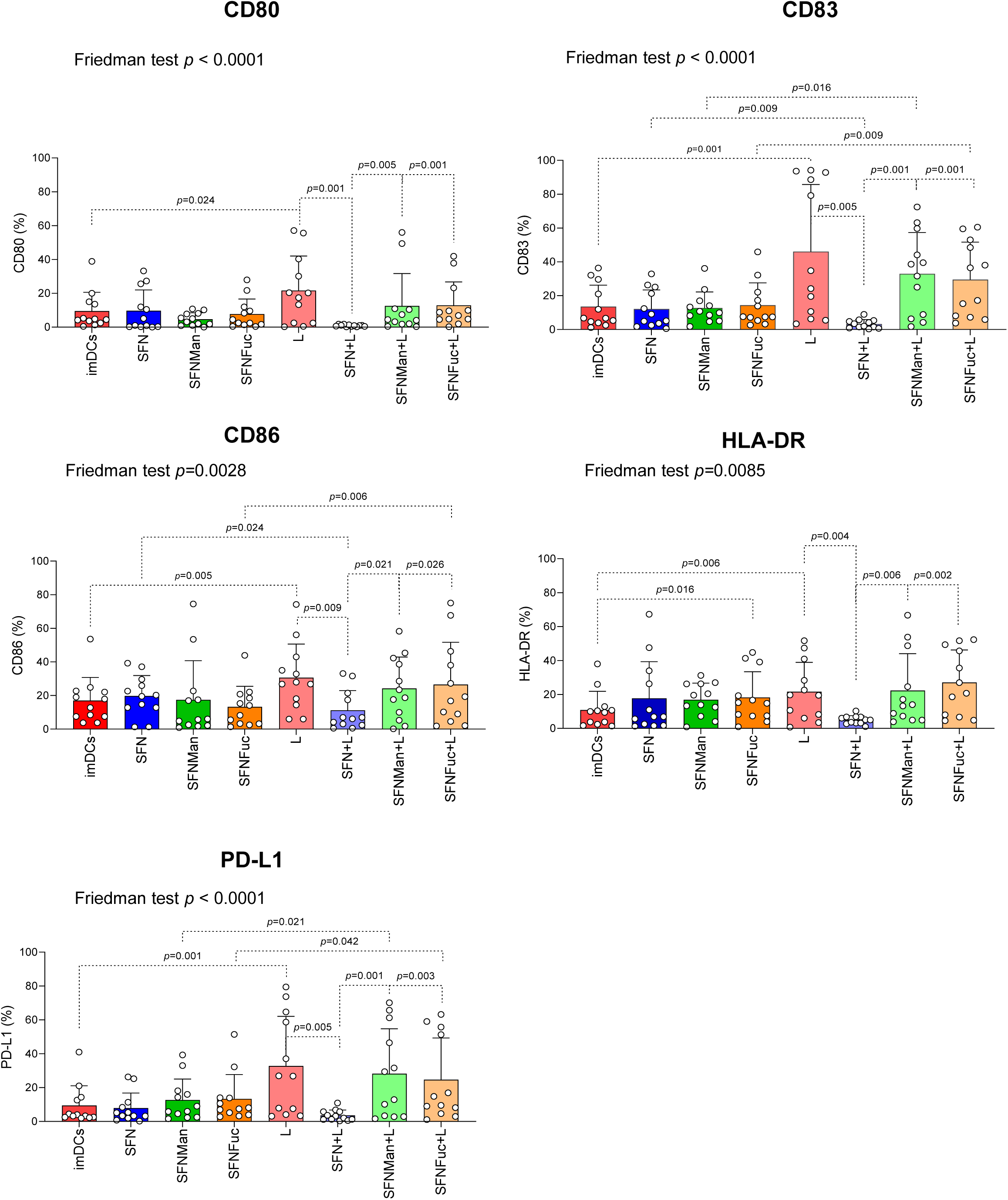
Changes in the expression of DCs surface molecule. The bars and symbols represent the mean (SD) of the percentages of expression of CD80, CD83, CD86, PD-L1 and HLA-DR on moDCs under different experimental conditions (N=12). imDC: immature dendritic cells. L: Lipopolysaccharide, LPS. SFN: Sulforaphane. SFNMan and SFNFuc: SFN-glycoconjugates with mannose or fucose. S+L: SFN incubated for 1 hour followed by LPS stimulation. SFNMan+ and SFNFuc+L: SFN-glycoconjugates incubated for 1 hour followed by LPS stimulation. The Friedman test was used to detect differences in related samples across multiple comparisons, indicating significant p-values. The Wilcoxon test was used for pairwise comparisons of related samples, indicating the exact p values.

Furthermore, to explore whether SFN and SFN-glycoconjugates could influence the expression of moDC regulatory molecules in the presence of LPS (to induce an inflammatory microenvironment), moDCs were preincubated with SFN and SFN-glycoconjugates for 1 hour, followed by LPS stimulation (Figure 2). A significant decrease in the expression of all markers was observed compared to moDCs pretreated with SFN (SFN+L) and LPS alone. However, irrespective of the carbohydrate in the SFN-glycoconjugates (SFNMan+L and SFNFuc+L), there was a significant upregulation of all markers compared to the effects of S+L. Notably, this increase in marker expression was significantly higher than the percentages observed in the absence of LPS for the same glycoconjugates, particularly for CD83 and PD-L1. Additionally, a significant increase in CD86 expression was also observed between SFNFuc+L *vs.* SFNFuc (Figure 2). However, no differences in the phenotypic state of the moDCs were observed, regardless of the carbohydrate present in the SFN-glycoconjugates.

### SFN-glycoconjugates reduce the pro-inflammatory cytokines

To evaluate cytokine production, supernatants were collected from the moDC maturation experiments. The production of pro-inflammatory cytokines IL-5 and TNF-α tended to decrease in moDCs pretreated with SFN and its glycoconjugates containing mannose and fucose, compared to LPS alone (Figure 3). This highlights the potential of SFN-glycoconjugates, considering carbohydrates as adjuvants, as they can enhance the protective effect of SFN solely against inflammatory stimuli such as LPS. However, this trend was not observed for the other pro-inflammatory cytokines, such as IFN-γ and IL-17 (Figure 3).

**Figure 3.**
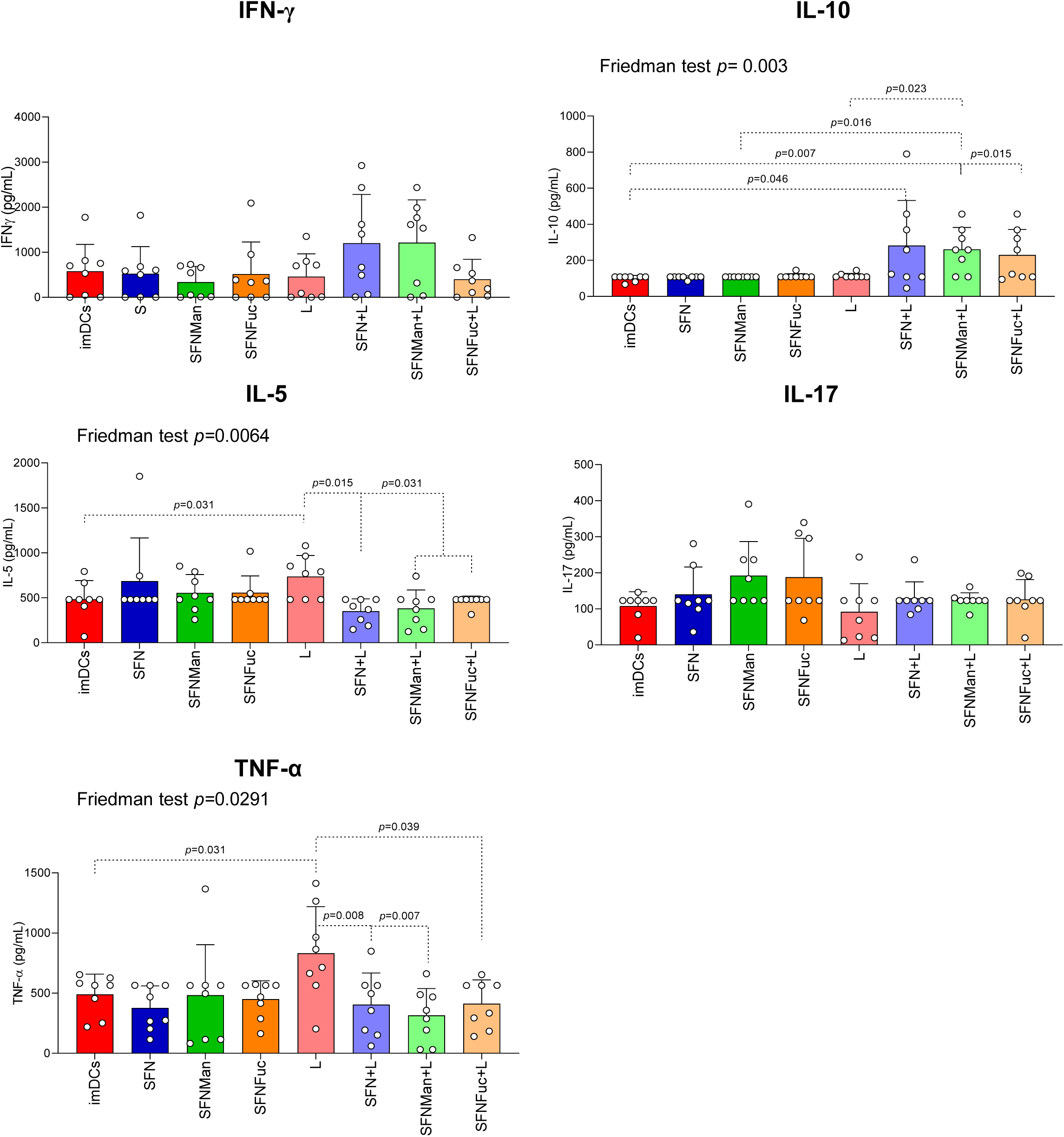
Changes in the cytokine production on DCs. The bars and symbols represent the mean (SD) of the concentration of each cytokine (pg/mL) (N=8) on moDCs under different experimental conditions, imDC: immature dendritic cells. L: Lipopolysaccharide, LPS. SFN: Sulforaphane. SFNMan and SFNFuc: SFN-glycoconjugates with mannose or fucose. S+L: SFN incubated for 1 hour followed by LPS stimulation. SFNMan+ and SFNFuc+L: SFN-glycoconjugates incubated for 1 hour followed by LPS stimulation. The Friedman test was used to detect differences in related samples across multiple comparisons, indicating significant p-values. The Wilcoxon test was used for pairwise comparisons of related samples, indicating the exact p values.

Regarding the regulatory cytokine IL-10, a significant increase in production was observed in moDCs pretreated with SFN and SFN-glycoconjugates before LPS restimulation (Figure 3).

### SFN-glycoconjugates induce changes in the proliferative response

moDCs were pre-incubated with SFN and SFN-glycoconjugates in the absence and presence of LPS (100 ng/mL), washed, and then used as APCs by co-culturing them with autologous lymphocytes. Proliferation of CD3^+^, CD3^+^CD4^+^FOXP3^+^T-, and CD3^-^CD19^+^B-cells was assessed.

SFNMan significantly increased CD3^+^CD4^+^FOXP3^+^T-cell proliferation under LPS stimulation compared to unstimulated cells. Furthermore, SFNMan+L induced significantly higher proliferation of CD3^+^CD4^+^FOXP3^+^T-cells compared to SFN+L. However, only SFN pretreatment led to a significant increase in IL-10 production by proliferated CD3^+^CD4^+^FOXP3^+^T-cells following LPS stimulation (Figure 4). This protective effect of SFN and SFNMan-conjugate after LPS restimulation also significantly increased CD3^-^ CD19^+^B-cell proliferation compared to SFN and SFNMan, respectively. Notably, SFNMan enhanced CD3^-^CD19^+^B-cell proliferation compared to unstimulated cells, suggesting that the nature of the SFN-glycoconjugates can influence the immune response (Figure 4). Conversely, SFNFuc did not promote a proliferative immune response, emphasizing the importance of carbohydrate composition in determining cellular outcomes (Figure 4).

**Figure 4:**
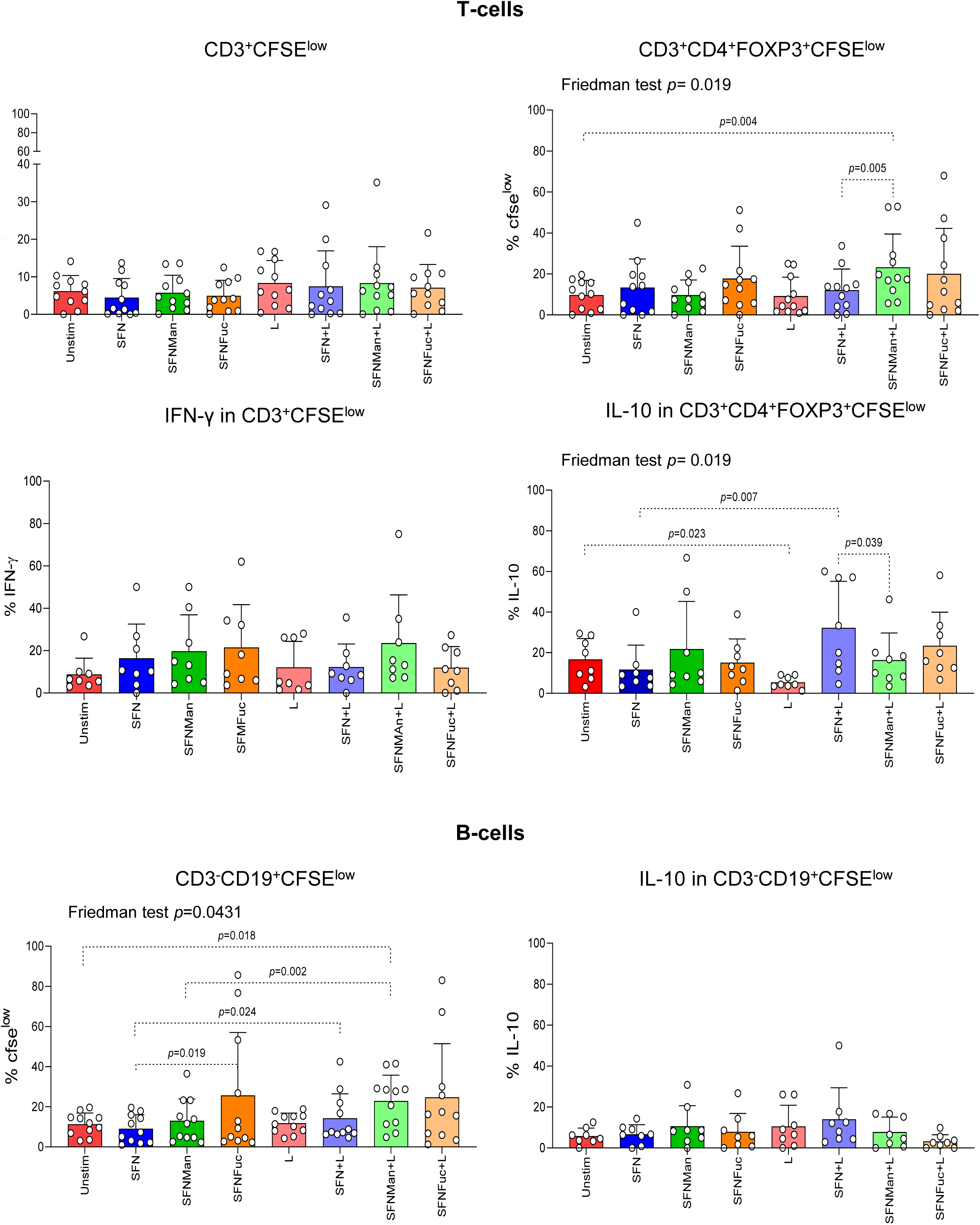
Proliferative T and B response induced by the SFN-glycoconjugates pretreatment. The bars and symbols represent the mean and standard deviation of the different percentages of CFSE_low_ for T-, B- and Treg-cells, and the percentages of IFN-γ and IL-10 levels in proliferating T-, Treg- and B-cells under different experimental conditions. (N=8-11). Unstim: Unstimulated cells. L: Lipopolysaccharide, LPS. SFN: Sulforaphane. SFNMan and SFNFuc: SFN-glycoconjugates with mannose or fucose. S+L: SFN incubated for 1 hour followed by LPS stimulation. SFNMan+ and SFNFuc+L: SFN-glycoconjugates incubated for 1 hour followed by LPS stimulation. CFSE: 5,6-carboxyfluorescein diacetate N-succinimidyl ester. The Friedman test was used to detect differences in related samples across multiple comparisons, indicating significant p-values. The Wilcoxon test was used for pairwise comparisons of related samples, indicating the exact p values.

### Effect of SFN-glycoconjugates on moDCs activation through inhibition of p65 NF-κB

The study also aimed to determine whether pretreatment of moDCs with MG132 altered the expression of CD80, CD83, CD86, HLA-DR, and PD-L1 following LPS stimulation, given that SFN-glycoconjugates suppressed p65 NF-κB, when challenged with LPS. To achieve this, moDCs were pretreated with SFN-glycoconjugates and MG132 for 1 hour before LPS addition, and this condition was maintained for 48 hours.

In the absence of an inflammatory microenvironment, significant differences in CD86 expression were observed between moDCs treated with MG132 *vs*. SNF, *vs.* SFNMan and *vs.* immature moDCs, including SFNFuc *vs.* MG132 (Figure SI2A).

In addition, our findings revealed that pretreating moDCs with SFN-glycoconjugates for 1 hour followed by LPS significantly increased the expression of CD83, CD86 and HLA-DR compared to MG+L. Additionally, SFN-glycoconjugates+L significantly increased CD80 and CD83 expression compared to SFN+L, while no differences were observed for PD-L1 expression (Figure 5A).

**Figure 5:**
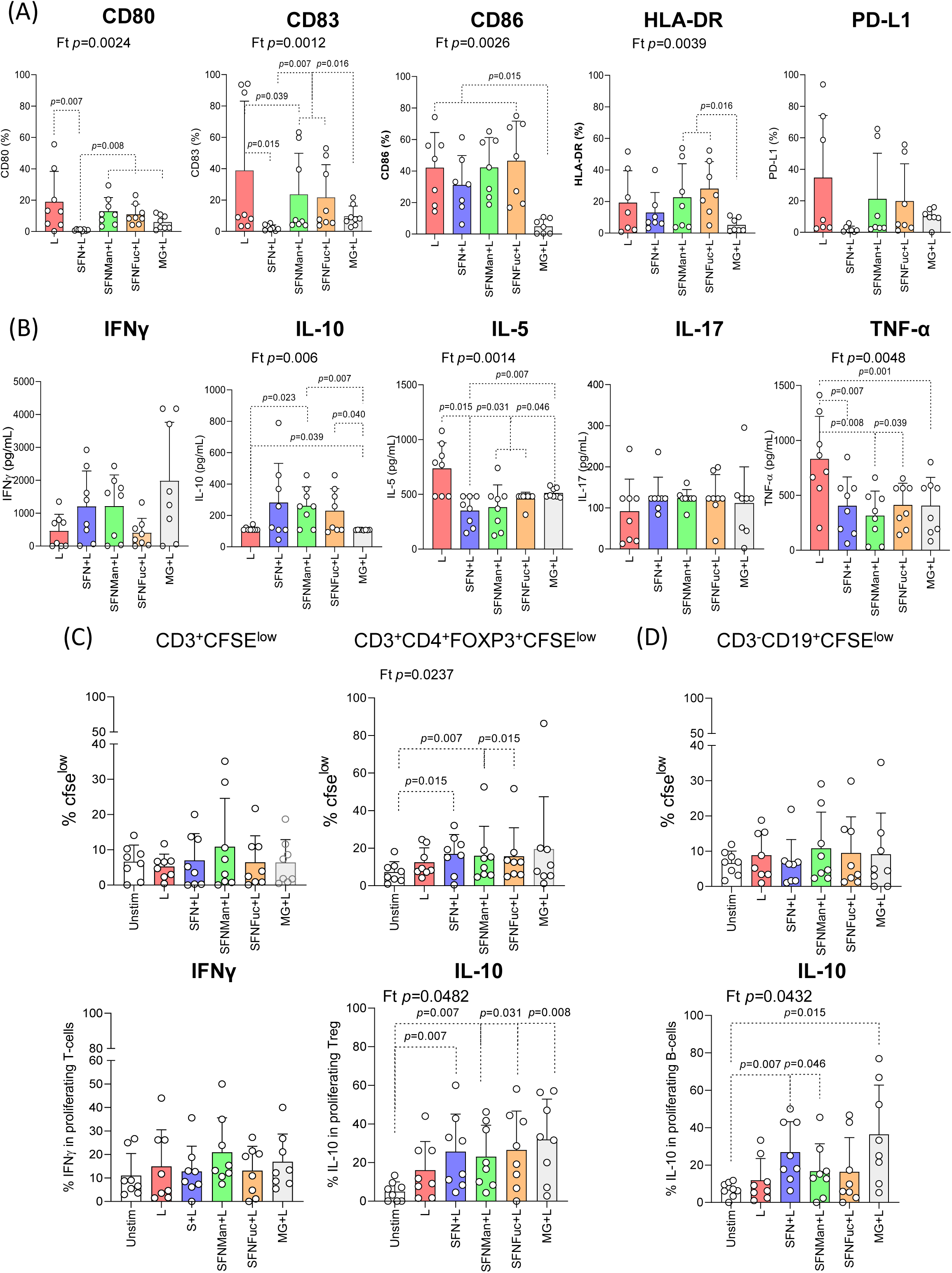
The effect of SFN-glycoconjugates is more potent than inhibition of p65 NF-κB. The bars and symbols represent the mean (SD) of the different percentages of expression of **(A)** CD80, CD83, CD86, PD-L1 and HLA-DR, of **(B)** the concentration of each cytokine (pg/mL), of **(C)** CFSE^low^ for T-, and Treg-cells and the percentage of IFN-γ and IL-10 levels in proliferating T- and Treg-cells, and **(D)** of CFSE^low^ for B-cells and the percentage of IL-10 levels in proliferating B-cells. (N=8). CFSE: 5,6-carboxyfluorescein diacetate N-succinimidyl ester. imDCs: immature DCs. Unstim: Unstimulated cells. L: Lipopolysaccharide, LPS. S+L: SFN incubated for 1 hour followed by LPS stimulation. SFNMan+ and SFNFuc+L: SFN-glycoconjugates incubated for 1 hour followed by LPS stimulation. MG+L: MG132, blocker for p65 p65 NF-κB, incubated for 1 hour followed by LPS stimulation. The Friedman test (Ft) was used to detect differences in related samples across multiple comparisons, indicating significant p-values. The Wilcoxon test was used for pairwise comparisons of related samples, indicating the exact p values.

Only SFN-glycoconjugates pretreatment followed by LPS stimulation led to increased IL-10 production compared to MG+L. Regarding treatment effects, the insertion of mannose (SFNMan+L) significantly increased IL-10 compared to LPS alone (Figure 5B). Pretreatment with SFN and SFN-glycoconjugates, followed by LPS stimulation and MG+L, significantly reduced IL-5 and TNF-α levels compared to LPS (Figure 5B). Notably, IL-5 levels were markedly reduced in the presence of SFN and its conjugates (SFN+L and SFNMan+ or SFNFuc+L) compared to MG+L. However, our results did not reveal significant differences in cytokine levels in the absence of LPS restimulation (Figure SI2B).

No differences were observed in the proliferation of T- and B-cells in the absence of LPS (Figure SI2C-D). However, the absence of LPS affected IL-10 levels in proliferating B-cells, with significant increases noted in the presence of the p65 NF-κB inhibitor (MG132) compared to unstimulated cells and SFN-glycoconjugates, as indicated in Figure SI2D.

We also analyzed the effect of the MG132 blocker on cellular proliferation. Blocking the p65 NF-κB pathway followed by LPS stimulation resulted in similar proliferation levels of Treg-cells stimulated by SFN+L, SFNMan+L and SFNFuc+L, compared to unstimulated DCs (Figure 5C). Regarding IL-10 levels, blocking p65 NF-κB followed by LPS stimulation led to an increase in IL-10 levels comparable to those observed under SFN+L, SFNMan+L and SFNFuc+L stimulation. This suggests that SFN-glycoconjugate pretreatment effectively regulates cell proliferation and enhances IL-10 production, comparable to the effects observed when blocking the p65 NF-κB pathway. Interestingly, blocking p65 NF-κB significantly increased IL-10 levels in the presence of SFN+L and SFNMan+L compared to unstimulated cells in proliferating B-cells (Figure 5D), achieving similar IL-10 production levels to those observed with MG+L pretreatment.

These findings underscore the potential of glycosylated SFN forms as pretreatment for therapeutic applications, enhancing their immunomodulatory effects without increasing cytotoxicity.

### The protective effect of SFN-glycoconjugates is independent of MAPKs inhibition

Given that SFN-glycoconjugates did not suppress MAPK (p38 MAPK and JNK) expression, we sought to confirm that their regulatory effects are independent of this signaling pathway. Phenotypic changes in moDCs were analyzed as previously described, and the proliferative responses of T-, Treg-, and B-cells, as well as their production of pro-inflammatory and regulatory (IL-10) cytokines, were examined in the presence and absence of specific blockers (SB203580 and SP600125).

Our results demonstrated that SFNMan induced a significant increase in the expression of CD83 and CD86 compared to the inhibitory effect of p38 MAPK after LPS restimulation (SB+L) (Figure SI3A). However, for SFNFuc, a significant increase in CD80 expression was observed when the p38 MAPK signaling pathway was inhibited after LPS restimulation (Figure SI3A). These results suggest that SFN-glycoconjugates exert a strong protective effect on cell activation, which is not replicated when the p38 MAPK signaling pathway is suppressed.

Moreover, SFNFuc pretreatment significantly reduced T-cell proliferation and IFN-γ production compared to MAPK blockade (SB203580 and SP600125) following LPS restimulation. This effect was not observed in Treg- and B-cell proliferation, consistent with the lack of IL-10 production (Figure SI3B). Thus, the regulatory effects of SFN-glycoconjugates appear independent of MAPK inhibition.

## DISCUSSION

The use of nano-delivery systems, specifically carbohydrate-functionalized nutraceuticals, in biomedical applications has grown significantly in recent years (Gowd et al., 2022; Pramudya & Chung, 2019). These systems have been proposed as therapies for various pathologies, including inflammatory disorders and cancer, by modulating the inflammatory response (Mukhtar et al., 2020). Carbohydrates within these systems act as adjuvants, eliciting specific immune responses (Chowdhury, Toth, & Stephenson, 2022). Previous studies have suggested that carbohydrates such as mannose and fucose, acting as adjuvants, enhance uptake by moDCs and induce maturation and lymphocyte proliferation, particularly in food-allergic patients (Palomares et al., 2022; Palomares et al., 2019). Additionally, the modulation of macrophage M1/M2 polarization using carbohydrate-functionalized polymeric nanoparticles has been explored (Andrade, Reis, Costas, Lima, & Reis, 2020), and the multivalent glycosylation of gold nanoclusters has been showed to promote increased uptake through endocytic pathways in moDCs (Le Guevel et al., 2015).

Inspired by these findings, we synthetized two glycoconjugates incorporating sulforaphane by coupling an amino-derivative of sulforaphane with either a mannose- or fucose-configured carbohydrate bearing an isothiocyanate moiety.

Recent studies emphasize that SFN-nanocarries, such as nanoencapsulated SFN in membrane vesicles, significantly enhance SFN’s anti-inflammatory effects in *in vitro* human macrophage models, thereby improving its therapeutic efficacy (Mohanty, Sahoo, Konkimalla, Pal, & Si, 2021; Yepes-Molina et al., 2022). These advancements address the limitations associated with SFN’s low bioavailability (Ruhee & Suzuki, 2020). In this regard, our study doubled the concentration of SFN used in *in vitro* experiments without observing cytotoxic effects, suggesting that these glycoconjugates improve SFN’s efficacy in *in vitro* applications, potentially serving as a foundation for future clinical applications, as observed in other functionalized systems (Yepes-Molina & Carvajal, 2021).

At the molecular level, our findings demonstrated that both free SFN and SFN-glycoconjugates exert a protective effect against LPS-induced activation in moDCs. This inhibition is crucial, as NF-κB serves as a central regulator of inflammatory responses, with its inactivation leading to reduced pro-inflammatory cytokine (TNF-α) production. In agreement with these results, several *in vitro* studies have shown that SFN and its delivery nanosystems mitigate inflammatory responses (Mohanty et al., 2021; Yepes-Molina & Carvajal, 2021). Interestingly, pre-treatment with SFN reduced the activation of the MAPK signaling pathway in LPS-stimulated moDCs, particularly through the downregulation of JNK expression. However, this effect was not observed with SFN-glycoconjugates, likely due to chemical differences associated with glycosylation. These observations suggest that SFN glycosylation may facilitate SFN entry via interactions with C-type lectin receptors (CLRS), such as DC-specific intercellular adhesion molecule-3-Grabbing non-integrin (DC-SIGN) on moDCs, thereby inhibiting NF-κB signalling pathways (Lin et al., 2015) and triggering cross-activation with the MAPKs, as reported in gastric epithelial cells (Y. Chen, Huang, & Xu, 2020).

This anti-inflammatory effect was also evident immunologically. Free SFN had the ability to downregulate the phenotypic state of the moDCs towards an immature state (reducing the expression of CD80, CD83, CD86, HLA-DR and PD-L1) under conditions of chronic inflammation, consistent with our previous findings (Fernandez-Prades et al., 2023). In contrast, pre-treatment with SFN-glycoconjugates followed by LPS stimulation significantly upregulated these markers compared to treatment with free SFN. This suggests that SFN-glycoconjugates, irrespective of the carbohydrate moiety, promote complete moDC maturation and activation. These effects were accompanied by significantly increased in IL-10 production, along with low levels of IL-5 and TNF-α (Kuipers et al., 2020), facilitating interactions with other immune cells (B-cells) and driving T-cell polarization (Busold et al., 2020).

These results suggest that SFN-glycoconjugates stimulate the maturation of moDCs by inducing phenotypic changes distinct from those elicited by free SFN, presumably through interactions with CLRs on moDCs. Furthermore, our findings demonstrate that pre-treatment with SFN-glycoconjugates induces specific T- and B-cell proliferation characterized by a regulatory response pattern, including Treg proliferation, as reported previously (Chappell et al., 2014). Such interactions between CLRs and SFN-glycoconjugates could be instrumental in promoting tolerance rather than reactivity. Therefore, the chemical nature of these glycoconjugates plays a pivotal role in modulating immunological and molecular responses when compared to free SFN. The use of glycosystems significantly enhances human DC targeting, likely due to improved diffusion and binding to target cells (Le Guevel et al., 2015; Martinez-Bailen, Rojo, & Ramos-Soriano, 2023).

Our results also revealed that pretreating moDCs with SFN-glycoconjugates, followed by LPS stimulation, led to a significant increase in the expression of CD83, CD86, and HLA-DR compared to the effects of specific p65 NF-κB blockade (via MG132) followed by LPS stimulation. This indicates that these glycoconjugates exert a robust effect on moDC maturation, resulting in increased production of anti-inflammatory cytokines (IL-10) and Treg cell activation. The potent immunomodulatory effects of SFN-glycoconjugates appear to stem from their inhibition of the p65 NF-κB pathway, akin to the mechanisms observed with other nanoformulations associated with functional foods, such as resveratrol (Sharifi-Rad et al., 2021). Notably, resveratrol-loaded β-lactoglobulin nanospheres have been shown to significantly increase IL-10 expression, thereby attenuating inflammation (Gowd et al., 2022; Pujara et al., 2021). Accordingly, the inhibition of this pathway by SFN-glycoconjugates reduces inflammation while fostering an immune response characterized by a regulatory profile.

In contrast, a different pattern was observed with specific MAPK blockers (SB203580 and SP600125). Pre-treatment with SFN-glycoconjugates showed no significant effects on inflammatory or regulatory responses in immune cells exposed to an inflammatory environment. This outcome could be attributed to chemical modifications associated with glycosylation, which might modulate inflammatory and immunological responses through mechanisms independent of the MAPK signaling pathway in DCs under inflammatory conditions. Consistently, previous studies have demonstrated that selenomethionine, an antioxidant, can regulate inflammation in intestinal epithelial cells independently of MAPK pathway regulation (Huang et al., 2024).

Despite certain limitations of our study, such as the need for further investigation into the interactions of these glycoconjugates with CLRs to elucidate their internalization in moDCs and their potential clinical applications, our findings underscore that SFN glycoconjugates are recognized by immune cells and elicit a more potent immunological response than free SFN. This heightened response is achieved through diverse signalling pathways in moDCs exposed to an inflammatory environment. SFN-glycoconjugates exhibit significant potential as vehicles for immunoregulatory and anti-inflammatory therapies, owing to their specific chemical properties, which facilitate complete moDC maturation and orchestrate a distinct regulatory proliferative response. These findings align with previous research conducted by our group (Palomares et al., 2022; Palomares et al., 2019), further emphasizing the potential of SFN-glycoconjugates as promising candidates for the development of new molecular therapeutic modalities targeting immune-related diseases characterized by inflammatory components, such as ulcerative colitis.

## MATERIALS AND METHODS

### Reagents and Tools table

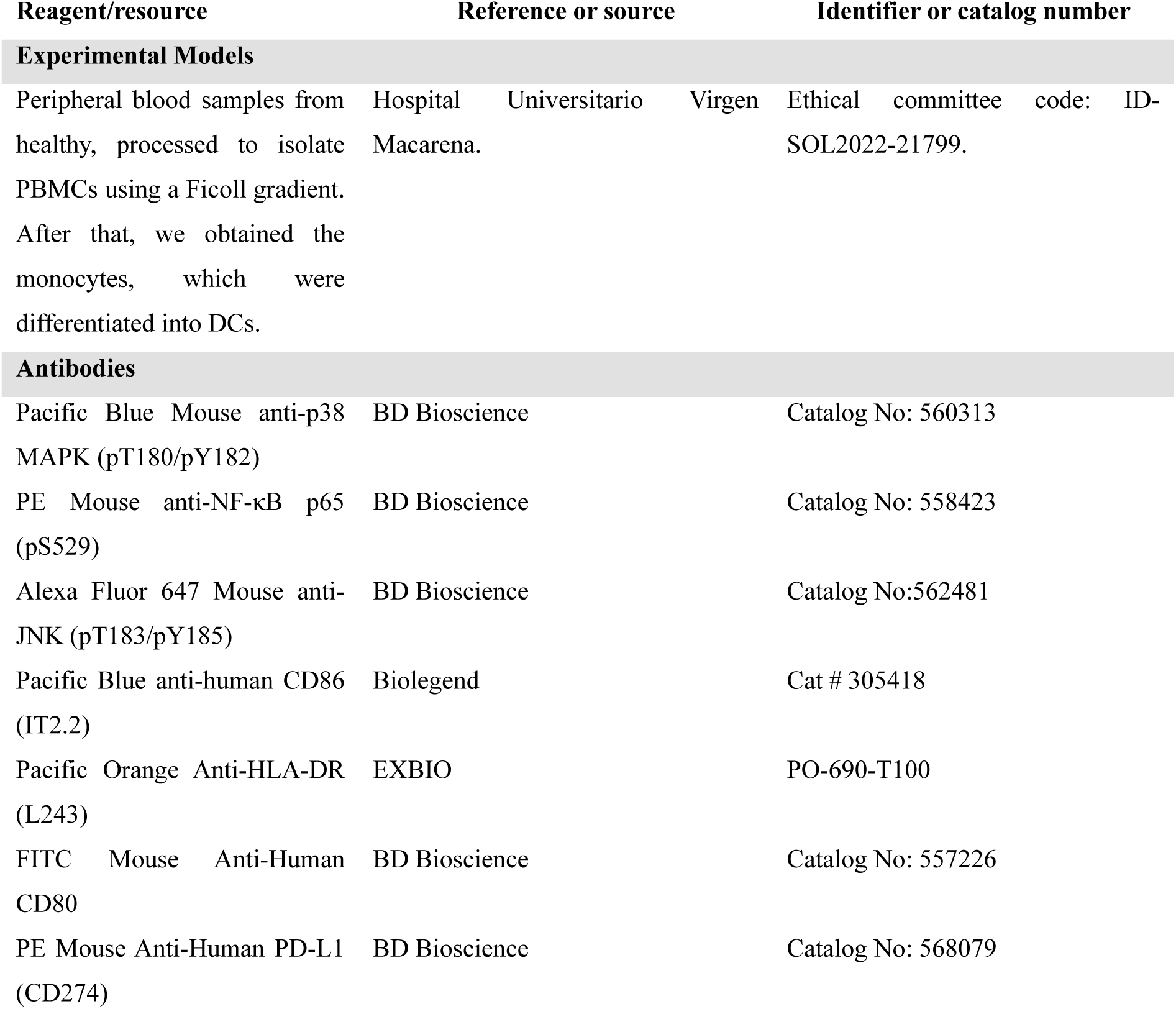
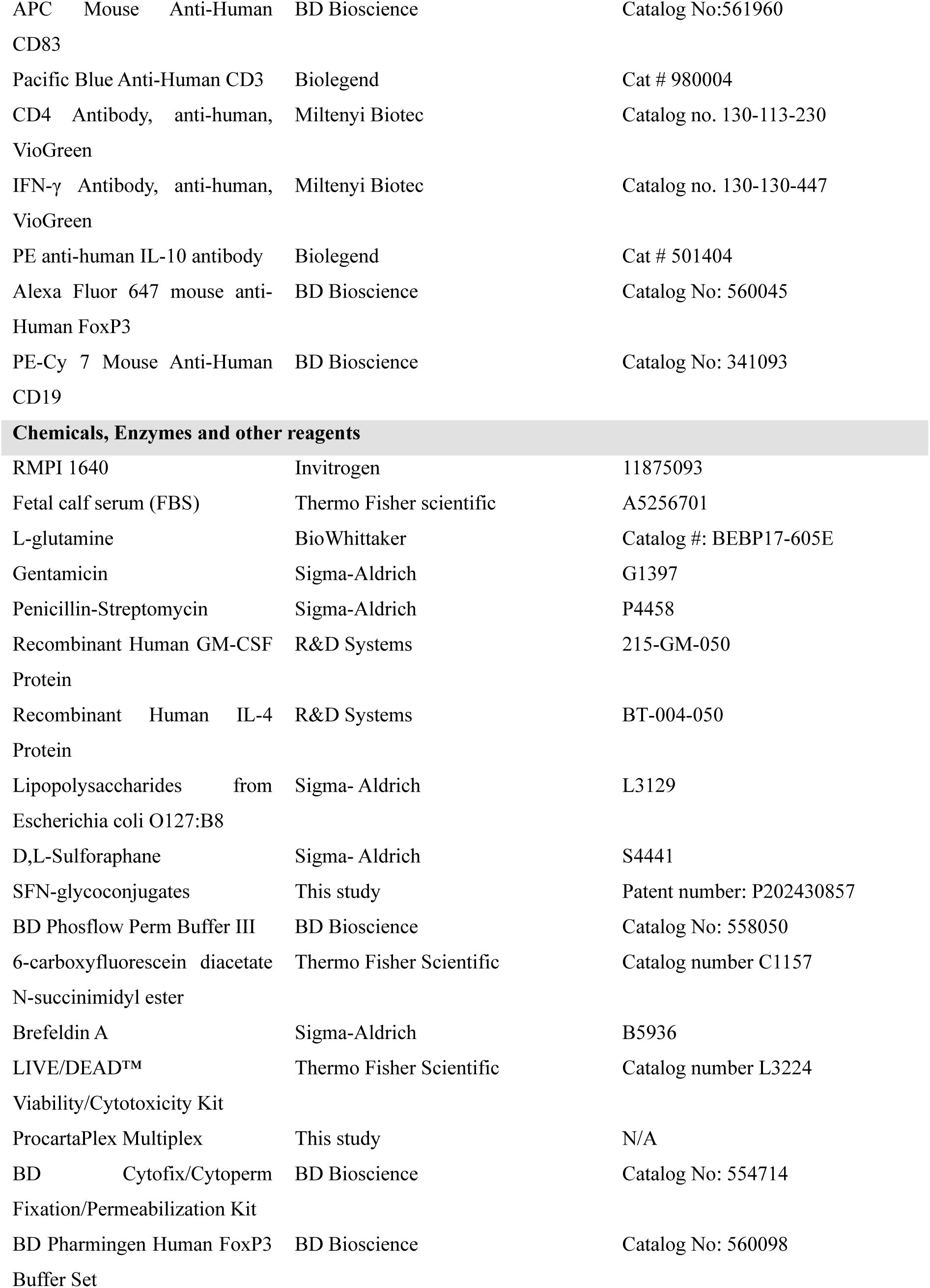
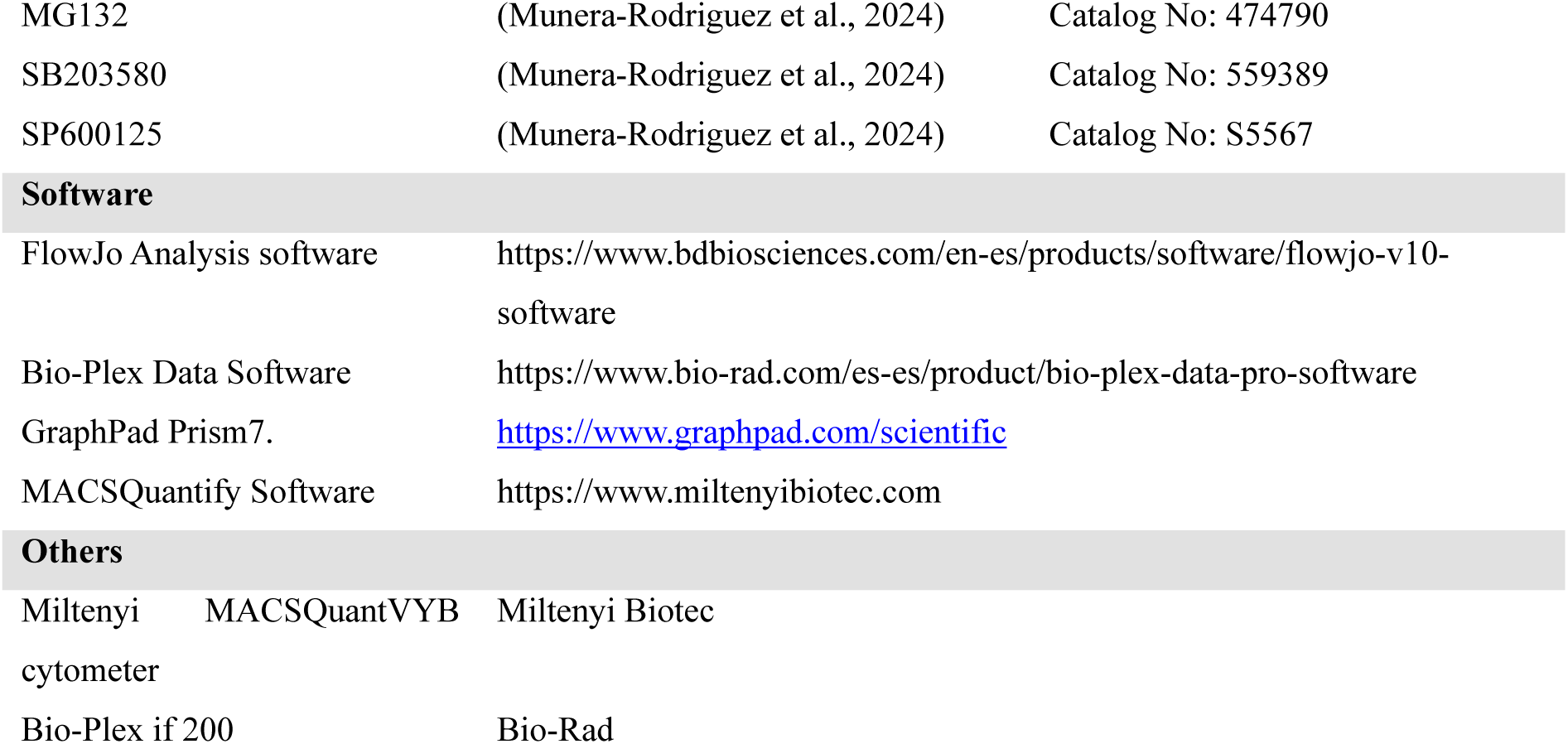

## Methods and Protocols

### Synthesis of SFN-glycoconjugates

The synthesis of SFN-glycoconjugates was performed through coupling reactions between an amino-derivative of SFN and a manno-or fuco-configured carbohydrate containing an isothiocyanate moiety (see supporting information for further experimental details).

### Study subjects

Biological samples of peripheral blood were sourced from healthy individuals aged 18 years or older who provided informed consent. Study participants had no recent history of immunoregulatory drug use or immunological disorders. Sample collection was approved by the Research Ethics Committee of Hospitales Universitarios Virgen Macarena, Virgen del Rocío under code: ID-SOL2022-21799.

### Sample Collection and Storing

Sample collection involved venipuncture to procure peripheral blood, from which peripheral blood mononuclear cells (PBMCs) were isolated using a Ficoll gradient. The PBMCs were subsequently cryopreserved in liquid nitrogen. All procedures followed standard protocols established by the HUVM-IBiS Biobank, which operates under the Andalusian Public Health System Biobank framework.

### Monocyte derived DCs (moDCs)

Monocyte-derived DCs (moDCs) were cultured in RMPI 1640 (Invitrogen, Gibco) supplemented with 10% fetal calf serum (FBS), 2 mM L-glutamine (BioWhittaker, Pittsburgh,PA), 5 mg/mL gentamicin and 50 ng/mL estreptomycin (both from Norton, Madrid, España). Differentiation was achieved by exposing the cells to 200 ng/mL GM-CSF and 100 ng/mL IL-4 (both from R&D Systems Inc, Minneapolis, MN), the cultured was maintained at 5% CO_2_ and 37°C.

### MoDCs cultures

To identify modifications induced by glycoconjugates in moDCs, the cultures were subjected to various experimental conditions. MoDCs (10 x 10^5^) were incubated in 96-well plates (Thermo-Fisher Scientific Waltham, MA) for 10 minutes at 37 °C and 5% CO_2_.

Experimental conditions included, mannose SFN-conjugate (SFNMan), fucose SFN-conjugate (SFNFuc) both 25 µM, lipopolysaccharides (LPS) from Escherichia coli 0127: B8 100 ng/mL as a positive control to induce chronic inflammation (Sigma-Aldrich, St. Louis, MO, USA), and SFN 10 µM (Sigma-Aldrich), and unstimulated moDCs serving as the negative control.

The protective properties and modifications induced by the SFN-glycoconjugates were examined by preincubating moDCs with SFN-glycoconjugates for 1 hour prior to LPS addition, thereby inducing a strong chronic inflammatory response. To compare their outcomes, the SFN was also incubated for 1 hour with LPS to generate chronic inflammation.

### Signaling Assay

MoDCs (7.5 × 10^5^) were grown in 96-well plates at 37°C and 5% CO_2_ for 10 minutes. For effective intracellular staining of p65 NF-κB and MAPKs, the moDCs were fixed with 1% paraformaldehyde and permeabilized using BD Phosflow Perm Buffer III (Becton, Dickinson, (BD) Franklin Lakes, NJ). Specific monoclonal antibodies (MoAbs) were utilized for this purpose (Reagents and Tools table). Flow cytometry was employed to assess changes in signaling pathways, measured using a Miltenyi MACSQuantVYB cytometer (Miltenyi Biotec, North Rhine-Westphalia, Germany) and analyzed in FlowJo software (BD). Results were presented as percentages of marker expression.

### Maturation Assay

MoDCs (10 × 10^5^) were cultivated in 96-well plates at 37°C and 5% CO_2_ after 48 hours. As previously stated, various experimental conditions were tested. To evaluate the effects of SFN-glycoconjugates, moDCs were preincubated with the glycoconjugates for 1 hour before the addition of LPS. The cells were stained with fluorochrome-labeled moAbs (Reagents and Tools table) to evaluate their maturation. Flow cytometry was used to analyze the expression levels of maturation markers, including CD80, CD83, CD86, HLA-DR, and PD-L1. Cell vitality was assessed using the LIVE/DEAD viability kit (Thermo Fisher Scientific). Flow cytometry data were collected with a Miltenyi MACSQuantVYB cytometer (Miltenyi Biotec) and analyzed using FlowJo software (BD). Results were expressed as the frequency of expression for each surface marker on the moDCs.

### Cytokine determination

Cytokine production (IFN-γ, IL-17, IL-5, TNF-α and IL-10) was quantified using a human ProcartaPlex Multiplex (Thermo Fisher Scientific). After 48 hours, supernatants from moDC cultures were collected. Samples (80 µL) and standards were incubated with a magnetic bead mix overnight at 4 °C. Subsequently, biotinylated antibodies were added and incubated for 300 minutes at room temperature (RT). Streptavidin was then added and incubated for another 300 minutes at RT. Following each incubation, the plate was washed and prepared for detection using Bio-Plex if 200 (Bio-Rad, Laboratories, Inc Hercules, CA, USA) with a reading buffer. Data analysis was performed using Bio-Plex Data Analysis Software (Bio-Rad Bio-Rad), and results were expressed as the concentration (pg/mL) of each cytokine.

### Specific proliferative response

As previously described, autologous moDCs (1.5 x 10^3^) in 96-well plate were pre-stimulated with SFNMan, SFNFuc 25 µM and SFN in the presence or absence of LPS for 48 hours Proliferation analysis was performed with 5,6-carboxyfluorescein diacetate N-succinimidyl ester (CFSE) (Thermo Fisher Scientific), utilizing 1.5 × 10^5^ / mL prelabeled CD14-cells. These cells were exposed to moDCs that had been pre-stimulated with different experimental combinations (10:1 ratio) in a final volume of 250 μL of the whole medium for 6 days at 37 °C and 5% CO_2_.

Furthermore, CFSE was used to evaluate proliferative responses of T- and B-cells, in addition to determining the lymphocyte subsets (CD3^+^T-, CD3^−^CD19^+^B-, and CD3^+^ CD4^+^FOXP3^+^T-cells, as Treg-cells). Detailed information on the fluorescent moAbs employed is available in the Reagents and Tools table.

The cytokine production capacity was confirmed by measuring IL-10 and IFN-γ levels. Brefeldin A (5 mg/mL) (Sigma-Aldrich) was added at a concentration of 1/1000 for 3 hours, followed by fixation with the Cytofix/CytoPerm Fixation/Permeabilization Solution Kit (BD). To mark regulatory T cells (Treg) a human FOXP3 buffer (BD) was used for Treg-cell staining. Results were expressed as the percentage of CFSE^low^ for each cell subpopulation under different experimental conditions, as well as the percentage of IL-10 and IFN-γ in proliferating T- Treg- and B-cells.

### Blocking Assay

To investigate the role of the p65 NF-κB and MAPK signalling pathways in the immune response induced by SFN and SFN-glycoconjugates in moDCs, blocking assays were conducted. MoDCs were pre-incubated with specific blockers for 1 hour at 37°C. These blockers included MG132 (1 µM), a potent inhibitor of p65 NF-κB activation (Tanimura, Nakazato, & Tanaka, 2021), SB203580 (10µM), known for its ability to inhibit p38 MAPK (Lin et al., 2015), and SP600125 (10µM), functioning as a JNK inhibitor, preventing the activation of inflammatory genes (Lin et al., 2015), regardless of the absence or presence of LPS.

Changes in the signalling pathways (p65 NF-κB and MAPK) were assessed at 10 minutes, while alterations in the maturation state were measured at 48 hours as previously described. Results were expressed as the frequency of expression for each surface marker on the moDCs.

Additionally, alterations in the proliferative response of various T- and B-cell subgroups were studied. Autologous moDCs were pre-incubated with the described blockers, followed by specific assays for the proliferative response, expressed as the percentage of CFSE^low^ for each cell subgroup under varying experimental conditions, along with the percentage of IL-10 and IFN-γ present in proliferating T-, Treg- and B cells.

### Statistical analysis

For variables that did not display a significant normal distribution, non-parametric tests were applied. Comparisons of related samples were conducted utilizing the Friedman test to identify discrepancies among experimental conditions in multiple comparisons. A p-value threshold of 0.05 was used to indicate significance. Upon identifying significant differences via the Friedman test, subsequent pairwise comparisons of related samples were carried out using the Wilcoxon. Significant differences were represented by p-values. Statistical analyses were conducted using GraphPad Prism7.

## Data Availability

Data presented in this work are available from the authors upon request.

## Acknowledgements

This work was supported by the Ministerio de Ciencia e Innovación (Grant PID2020-116460RB-100 funded by MCIN/AEI/10.13039/50110001103, grant PID2023-147334OB-I00 funded by MICIU/AEI/ 10.13039/501100011033 and, as appropriate, by “ERDF A way of making Europe”, by “ERDF/EU”, by the “European Union”, and by Ramon y Cajal program (RYC2021-031256-I) funded by MICIU/AEI/10.13039/501100011033 and, as appropriate, by “ESF Investing in your future”, by “ESF+” or by “European Union NextGenerationEU/PRTR”), by the “VII Plan Propio de Investigación y Transferencia” from the University of Seville (Project-2023/00000482), and by Asociación Universitaria Iberoamericana de Postgrado scholarship for the Medical Research: Clinical and Experimental Master.

We also thank CITIUS Universidad de Sevilla (MS and NMR services, including the flow cytometry services) and Edijs Jansons for technical assistance.

## Author contributions

A.M. M-R and C. L-C: Writing—original draft preparation, Methodology and Writing-Reviewing and Editing. M. M-B and A. T C: Writing—original draft preparation, Methodology and Writing-Reviewing and Editing. Resources. Formal analysis, Investigation. Project administration. Resources. S. L-E: Supervision. Writing-Reviewing and Editing. F. P: Writing—original draft preparation, Conceptualization, Supervision, Writing-Reviewing and Editing. Formal analysis, Investigation. Project administration. Resources.

All authors have read and agreed to the published version of the manuscript. The authors declare that the described work has not been previously published and is not under consideration for publication elsewhere.

